# BayesFactorFMRI: Implementing Bayesian second-level fMRI analysis with multiple comparison correction and Bayesian meta-analysis of fMRI images with multiprocessing

**DOI:** 10.1101/2020.08.21.261966

**Authors:** Hyemin Han

## Abstract

BayesFactorFMRI is a tool developed with R and Python to allow neuroimaging researchers to conduct Bayesian second-level analysis and Bayesian meta-analysis of fMRI image data with multiprocessing. This tool expedites computationally intensive Bayesian fMRI analysis through multiprocessing. Its GUI allows researchers who are not experts in computer programming to feasibly perform Bayesian fMRI analysis. BayesFactorFMRI is available via Zenodo and GitHub for download. It would be widely reused by neuroimaging researchers who intend to analyse their fMRI data with Bayesian analysis with better sensitivity compared with classical analysis while improving performance by distributing analysis tasks into multiple processors.

## Introduction

BayesFactorFMRI is a tool developed with R and Python to allow neuroimaging researchers to conduct Bayesian second-level analysis and Bayesian meta-analysis of fMRI data with multiprocessing (Han, 2019; Han and Park, 2019). Previous studies have shown that using Bayesian statistics in fMRI analysis can be a way to address limitations in classical analysis based on *p*-values (Wagenmakers, 2007; Han and Park, 2018). For example, a Bayes factor, which is one of the most frequently used statistical indicators in Bayesian inference, can show us to which extent observed data supports a hypothesis of interest unlike a *p*-value that merely shows the extremity of observed data given the hypothesis (Rouder *et al*., 2009; Han, Park and Thoma, 2018). Furthermore, Bayesian analysis is more robust against noise, which is a significant issue in fMRI research, compared with classical analysis even with a small sample size (Han, 2019).

However, there is a significant practical limitation in implementing Bayesian analysis in the context of neuroimaging. As the previous studies presented (Han, 2019), it would take up to ten hours to complete Bayesian analysis with fMRI data because up to nine hundred thousand voxels may have to be analysed included in each image file. In addition, given that Bayesian analysis is based on iterative observations of data, such iterative processes per se can also be time consuming.

In order to address the aforementioned limitation of Bayesian fMRI analysis, a multiprocessing implementation of the Bayesian method in BayesFactorFMRI was attempted. Given that each fMRI image file consists of up to nine hundred thousand voxels to be tested, it would be possible to improve performance by assigning the voxels to multiple processors. BayesFactorFMRI uses NIfTI (.nii) or ANALYZE (.img +.hdr) image files as input files. For Bayesian second-level analysis, contrast images files that are created from first-level analysis performed by other fMRI analysis tools (e.g., SPM, FSL, AFNI) can be used for inputs. Moreover, to determine how many voxels are tested, a mask file that specify voxels to be tested is also used as an input. For Bayesian meta-analysis, statistical images from previous fMRI studies that report *t*-or *z*-statistics in each voxel can be used. Output images are created in the NIfTI format. BayesFactorFMRI creates output images files that report calculated Bayes factor values for hypothesis testing and calculated effect size values in tested voxels.

In addition to the parallelization of Bayesian analysis, BayesFactorFMRI provides users with a graphical user interface (GUI; see Figure 1). Because BayesFactorFMRI consists of R and Python codes, users might not be able to conduct Bayesian analysis as they intend without any further guidance, if they do not have sufficient expertise in computer languages. Hence a simple GUI was created so that end users can perform Bayesian analysis by following directions. Directions about how to use the GUI for Bayesian second-level analysis are available at https://github.com/hyemin-han/BayesFactorFMRI/blob/master/HowTo_2nd.md, and those for Bayesian meta-analysis are available at https://github.com/hyemin-han/BayesFactorFMRI/blob/master/HowTo_meta.md.

**Figure 1.**
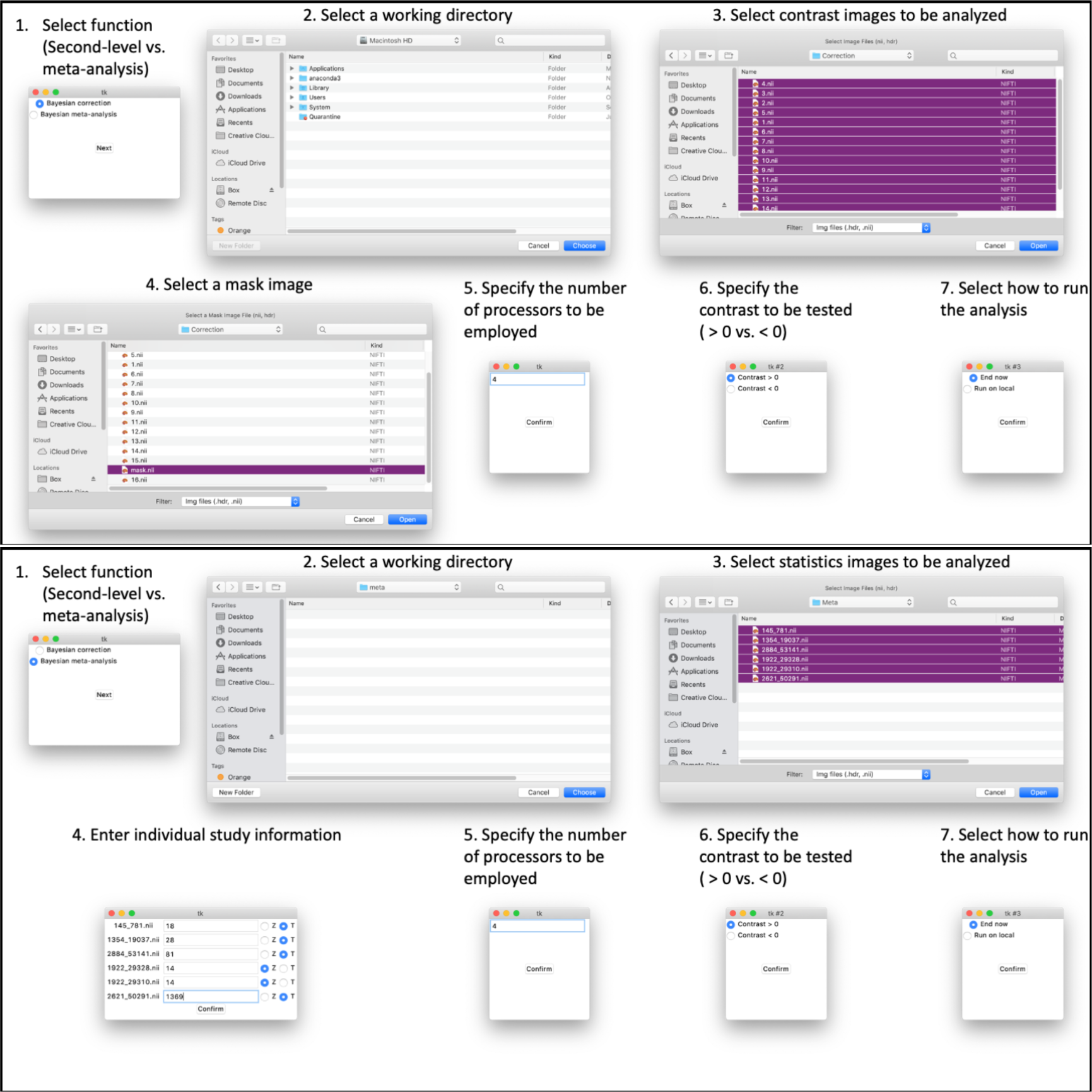
BayesFactorFMRI GUI. Top: GUI for Bayesian second-level analysis. Bottom: GUI for Bayesian meta-analysis.

The performances of Bayesian second-level analysis and meta-analysis were tested on the University of Alabama High-Performance Computing System (UAHPC; hardware and software specifications are described at https://oit.ua.edu/service/research/). In the case of Bayesian second-level analysis, the performances when 1, 2, 4, 8, 16, and 32 processors were employed were examined. For input images for the test, 16 NIfTI images that contain both true signals (activation foci extracted from) and the random noise were used. To create an original image with true signals, the activation foci that were reported in a previous neuroimaging study were utilized (Han, 2017). Figure 2 shows the original image with true signals (left) and noise-added image sample (middle). The analysis was repeated ten times per each processor condition and the mean elapsed time was calculated each time.

**Figure 2.**
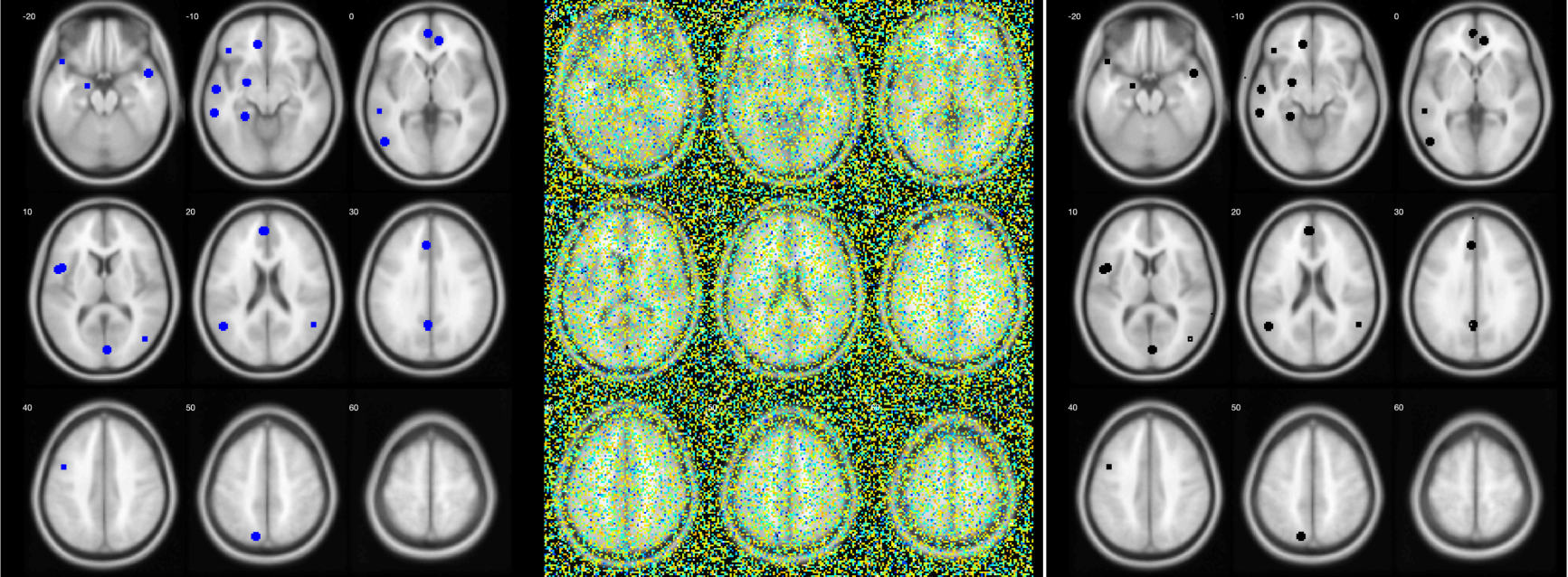
A tutorial example of Bayesian second-level analysis with BayesFactorFMRI. Left: The original image with true signals (blue: true positives). Middle: A sample of analysed images with original and noise signals. Right: The result of analysis when BFs.nii is thresholded with a Bayes factor threshold ≥ 3 (black: survived voxels).

In the case of Bayesian meta-analysis, 2, 4, 8, 16, 32, and 64 processors were used. To test the performance of Bayesian meta-analysis, six NIfTI images that were created in previous fMRI studies that examined the neural correlates of the working memory were analyzed. These six images were downloaded from NeuroVault (https://neurovault.org/), an open repository for statistics image files created by previous fMRI studies, by using a keyword, “working memory” (For the full list of the studies, refer to Table S1 in Han and Park (2019)). For each condition, the test was repeated ten times and the mean elapsed time was calculated each time. Figure 3 (left) shows these six meta-analysed images.

**Figure 3.**
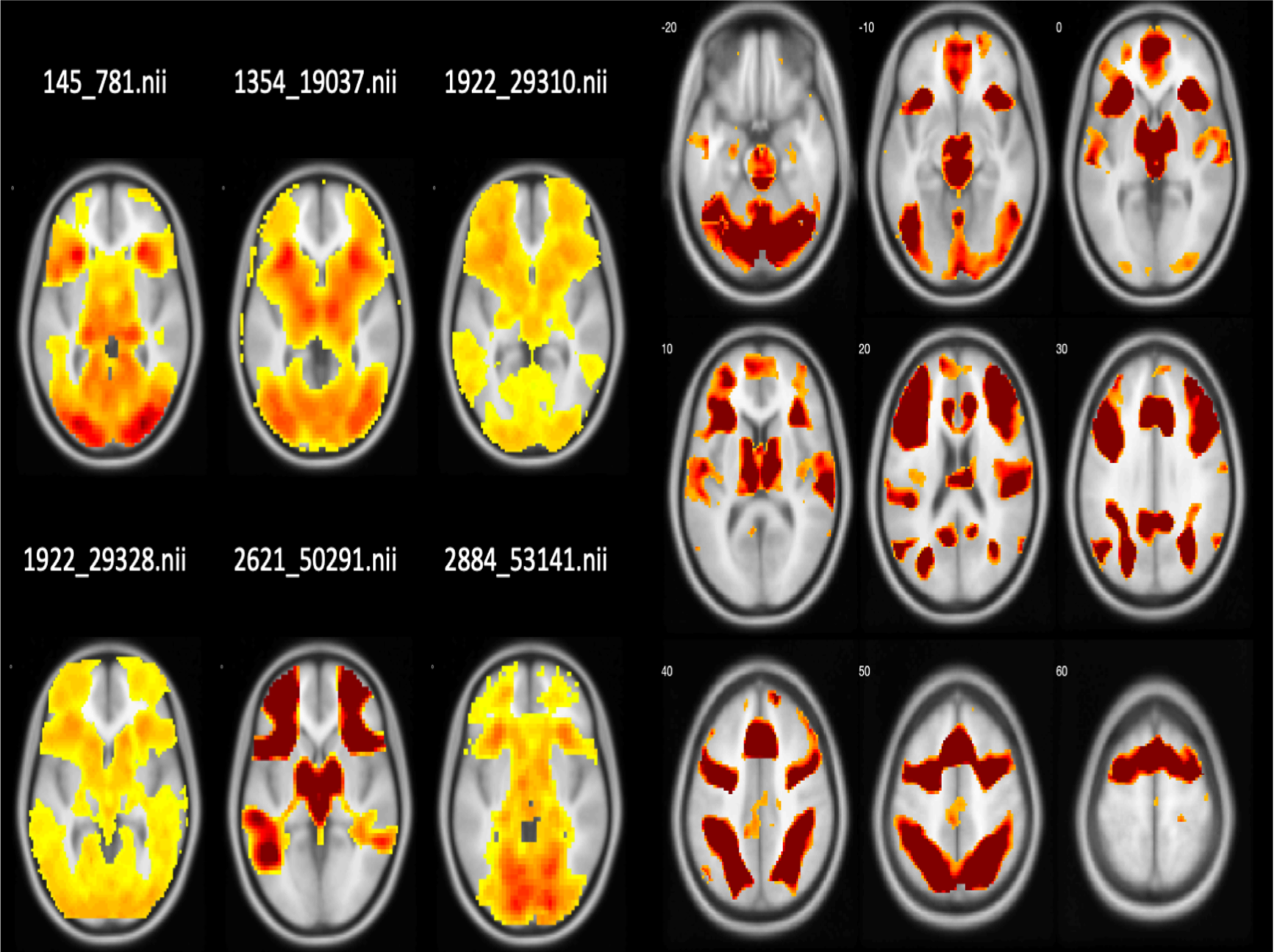
A tutorial example of Bayesian meta-analysis with BayesFactorFMRI. Left: Six statistics images that are used for meta-analysis. Right: The result of Bayesian meta-analysis when BFs.nii is thresholded with the Bayes Factor threshold ≥ 3. Only survived voxels are presented.

When the performances of Bayesian second-level analysis and meta-analysis with multiprocessing were tested, the increase of the number of employed processors resulted in the improved performance in terms of the decrease in the elapsed time. Figure 4 demonstrates the change in the elapsed time by the number of processors. As shown, the mean elapsed time decreased in a power scale as the number of employed processors increased.

**Figure 4.**
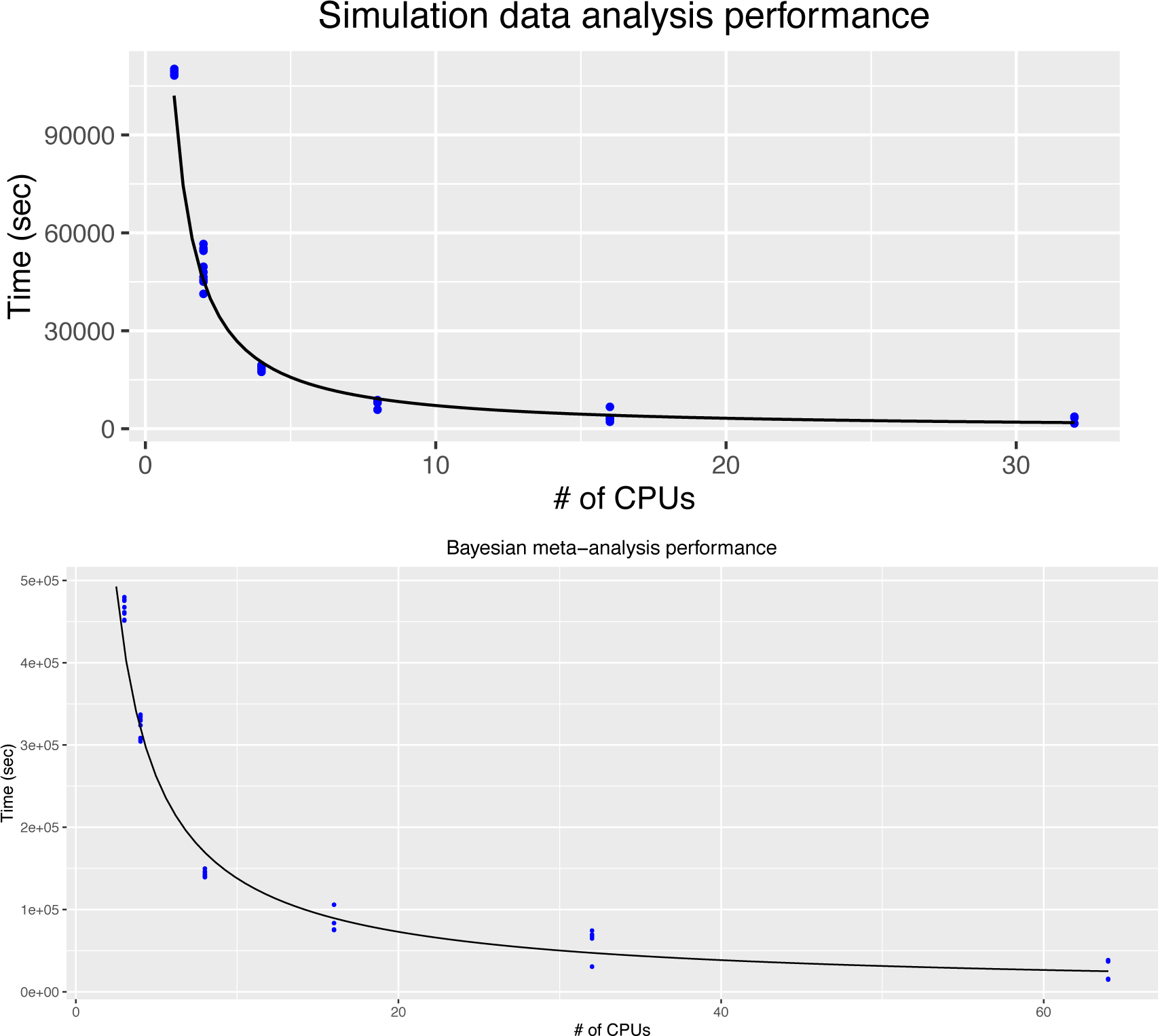
Changes in performance in terms of the elapsed time as a function of the number of employed processes. Top: Bayesian second-level analysis. Bottom: Bayesian meta-analysis.

### Implementation and architecture

The overall organization of BayesFactorFMRI is presented in Figure 5. Both Bayesian second-level analysis with multiple comparison correction and meta-analysis are performed with R codes based on BayesFactor package. In addition, custom Python codes are used to distribute voxels into different processors for multiprocessing and to present a GUI. The GUI of BayesFactorFMRI creates “run_this.py,” which calls the aforementioned codes for multiprocessing-applied Bayesian analysis. Following the user’s preference, run_this.py can be executed locally immediately after the closure of the GUI or can be uploaded to a computing cluster.

**Figure 5.**
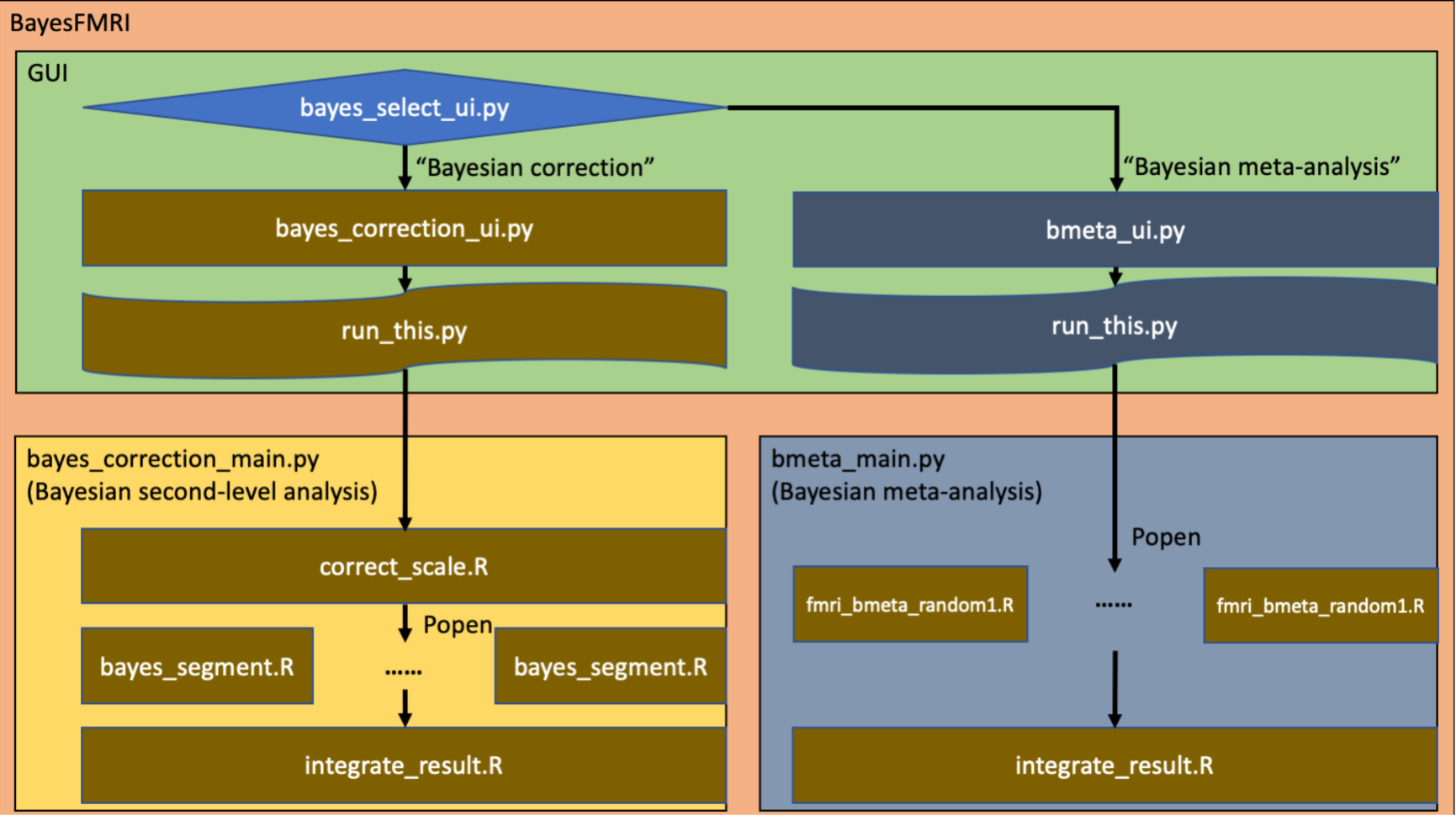
Organization of BayesFactorFMRI

When run_this.py is executed, a Python code that distributes voxels into different processors is called. In the case of Bayesian second-level analysis, “bayes_correction_main.py” is called, and in the case of Bayesian meta-analysis, “bmeta_main.py” is called. Both Python codes assign voxels into the designated number of processors so that workloads are evenly distributed to the processors. Then, R codes for the intended statistical analysis are called. By using “popen” in subprocess, the R codes are called multiple times according to the number of processors.

First, Bayesian second-level analysis is performed after adjusting the prior probability distribution to address multiple comparisons. This adjustment process occurs before multiprocessing is initiated. The prior probability, which follows the Cauchy distribution, is adjusted according to how many voxels are tested (performed by correct_scale.R). The number of voxels to be tested is specified by a mask image in NIfTI. Then, “popen” creates multiple processes to distribute voxels into different professors. Bayesian one-sample *t*-test is performed to examine whether there is a significant effect (e.g., a difference in neural activity) in each voxel (performed by Bayes_segment.R). Finally, integrate_result.R integrates the resultant Bayes Factor and median effect size value at each voxel in two NIfTI whole brain image files.

Second, Bayesian meta-analysis performed by performing random-effect meta-analysis of effect size values reported in previous studies in each voxel (fmri_bmeta_random1.R). At the beginning, “popen” calls multiple “fmri_bmeta_random1.R” to distribute voxels into the designated number of processors for multiprocessing. The effect size value in each voxel in each reported previous study is standardized. Then, Bayesian random-effect meta-analysis is performed at each voxel to examine whether there is a significant non-zero effect. Similar to the case of Bayesian second-level analysis, the analysis result in all voxels are integrated into whole-brain NIfTI image files. Three output images files, one reporting Bayes factors, one reporting mean effect size values, one reporting median effect size values, are created.

### Quality control

Both Bayesian second-level analysis and meta-analysis implemented in BayesFactorFMRI can be tested with tutorial datasets shared in GitHub (see https://github.com/hyemin-han/BayesFactorFMRI for further details). In the GitHub repository, further details regarding how to set options in the GUI are specified (refer to https://github.com/hyemin-han/BayesFactorFMRI/blob/master/README.md for general information, https://github.com/hyemin-han/BayesFactorFMRI/blob/master/HowTo_2nd.md for Bayesian second-level analysis, https://github.com/hyemin-han/BayesFactorFMRI/blob/master/HowTo_meta.md and for Bayesian meta-analysis). Following the directions, users can create “run_this.py” for either Bayesian second-level analysis or meta-analysis of the provided tutorial dataset. To initial the GUI, at the terminal, execute: python (or python3) bayes_select_ui.py. Once “run_this.py” is created, this code can be executed locally or uploaded to a cluster as per users’ preference. Whether the code is executed locally or uploaded to another place can be determined in the GUI.

Once “run_this.py” is executed, it calls R codes for either Bayesian second-level analysis or meta-analysis following the selected option. At the end of the analysis process, NIfTI files reporting analysis outcomes are created. When BayesFactorFMRI is executed successfully, users should be able to see two (in the case of Bayesian second-level analysis) or three (in the case of Bayesian meta-analysis) created NIfTI output files. In the case of Bayesian second-level analysis “BFs.nii” and “Ds.nii” are created. “BFs.nii” reports the resultant Bayes factor value and “Ds.nii” reports the median effect size value in Cohen’s *D* in each voxel. In the case of Bayesian meta-analysis, three output files, “BFs.nii,” “Medians.nii,” and “Means.nii,” are created. “BFs.nii” shows the resultant Bayes Factor value, “Medians.nii” does the median effect size value, and “Means.nii” does the mean effect size value in each voxel. For hypothesis testing (e.g., whether a significant non-zero effect exists in a voxel), users can open “BFs.nii” with a NIfTI viewer, such as xjView (Cui, Li and Song, 2015) with MATLAB, and perform thresholding (e.g., Bayes factor ≥ 3).

When the analysis is properly done with the provided tutorial dataset, if “BFs.nii” is thresholded with xjView plus MATLAB with a Bayes factor threshold ≥ 3, the result of Bayesian second-level analysis should be similar to Figure 2 (right). The result of Bayesian meta-analysis of the provided tutorial dataset after thresholding with the same program and Bayes Factor threshold should be similar to Figure 3 (right).

## (2) Availability

### Operating system

BayesFactorFMRI has been developed and tested on macOS Mojave (not tested on Catalina at this point) and Centos 7. Given that BayesFactorFMRI has been developed with R and Python, it can be executed on macOS, Linux, or Windows with compatible R and Python environments.

### Programming language

R (>= 3.5) and Python (>= 3.7.3; Python 3.8 is not recommended due to package-related issues at this point). Developed and tested on R 3.5 and Python 3.7.3.

### Dependencies

R: BayesFactor (developed with 0.9.12-4.2), metaBMA (developed with 0.6.1), oro.nifti (developed with 0.9.1) Python: tkinter (developed with 8.6), shutil, pandas (developed with 0.24.2), nibabel (developed with 2.4.1), rpy2 (developed with 3.2.2), numpy (developed with 1.16.2), nilearn (developed with 0.6.2), subprocess Specified directions about how to install required dependencies are available in https://github.com/hyemin-han/BayesFactorFMRI/blob/master/README.md

### List of contributors

Hyemin Han developed the software and created tutorials.

### Software location

***Archive*** (e.g. institutional repository, general repository) (required – please see instructions on journal website for depositing archive copy of software in a suitable repository)

***Name:*** Zenodo

***Persistent identifier:*** 10.5281/zenodo.3976338

***Licence:*** MIT License

***Publisher:*** Hyemin Han

***Version published:*** 1.0.0

***Date published:*** 08/08/20

**Code repository** (e.g. SourceForge, GitHub etc.) (required)

***Name:*** Github

***Identifier:*** https://github.com/hyemin-han/BayesFactorFMRI

***Licence:*** MIT License

***Date published:*** 08/08/20

***Language***

English

## (3) Reuse potential

BayesFactorFMRI is available via Zenodo and GitHub with tutorial datasets and directions. Given that it provides its potential users, neuroimaging researchers in particular, with the GUI, they will be able to perform Bayesian second-level analysis and meta-analysis with their fMRI dataset feasibly even without expertise in Python and R programming. Because BayesFactorFMRI is a tool to analyse functional neuroimaging data, it cannot analyse other types of images, such as anatomical brain images, spine or other soft medical/biological tissue images. In the current version, for Bayesian second-level analysis, only a simple one-sample t-test is supported. Overall, because BayesFactorFMRI is distributed via open repositories and a tutorial with testable image files and directions are available for users, it will be widely and straightforwardly reused by neuroimaging researchers who intend to apply Bayesian analysis with enhanced sensitivity and performance through multiprocessing.

## Acknowledgements

The author thanks Joonsuk Park and Ian M. McDonough for their comments on this work.

## Contact and support

Any bugs, errors, questions, or suggestions associated with BayesFactorFMRI can be submitted via the “Issues” tab in the GitHub repository. Furthermore, the author, Hyemin Han, can be contacted via email for support.

## Copyright Notice

Authors who publish with this journal agree to the following terms:

Authors retain copyright and grant the journal right of first publication with the work simultaneously licensed under a <u>Creative Commons Attribution License</u> that allows others to share the work with an acknowledgement of the work’s authorship and initial publication in this journal.

Authors are able to enter into separate, additional contractual arrangements for the non-exclusive distribution of the journal’s published version of the work (e.g., post it to an institutional repository or publish it in a book), with an acknowledgement of its initial publication in this journal.

By submitting this paper you agree to the terms of this Copyright Notice, which will apply to this submission if and when it is published by this journal.

